# Comparison of multiple cefiderocol susceptibility testing methods against genomic determinants of resistance in *bla*_NDM_ carbapenemase producing Enterobacterales

**DOI:** 10.64898/2026.01.27.701980

**Authors:** Caitlin Duggan, Daire Cantillon, David Lawrie, Timothy Neal, James Cruise, Fabrice E. Graf, Victoria Owen, Alice J. Fraser, Joseph M. Lewis, Charlotte Brookfield, Eva Heinz, Thomas Edwards

## Abstract

**Background:** Cefiderocol is a siderophore-conjugated cephalosporin antibiotic used to treat multi drug resistant Gram negative infections, including metallo-beta-lactamase producing Enterobacterales. Antimicrobial useage is guided by antimicrobial susceptibility testing (AST) which is hampered by differences between EUCAST and CLSI breakpoints, methodological challenges of AST, and lack of information on clinical outcome related to AST.

**Objectives:** This study assessed the agreement between AST methods under EUCAST and CLSI breakpoints in a collection of 57 *bla*_NDM_ producing Enterobacterales isolated from a UK hospital network.

**Methods:** All isolates, including *Klebsiella pneumoniae, Enterobacter hormaechei, Escherichia coli and Citrobacter freundii*, were whole-genome sequenced and tested with disk diffusion and MIC gradient test strip, and broth microdilution MICs were determined for a subset. Categorical agreement between methods was calculated using both EUCAST and CLSI breakpoints. Mutations and acquired resistance genes associated with cefiderocol resistance were identified and compared with AST results.

**Results:** The disk diffusion method, based on EUCAST interpretation, classified 94.7% of isolates as cefiderocol resistant and 5.3% as susceptible, with 22.8% within the Area of Technical Uncertainty. The CLSI breakpoint classified one isolate as resistant (1.8%) and 5.26% intermediate. Category agreement of broth microdilution and disk diffusion for E. coli using EUCAST guidelines was 38.5%. Mutations associated with cefiderocol resistance were highly prevalent and varied between species.

**Conclusions:** The discordant EUCAST and CLSI breakpoint values provided have large impacts on the classification of isolates susceptibility to cefiderocol, which will impact global cefiderocol usage and surveillance of resistance, further complicated by poor agreement between AST methods.

## Introduction

Antimicrobial resistance (AMR) is a major risk to global public health, with an estimated 1.27 million deaths directly attributed to bacterial AMR annually [1]. [2]Carbapenemase producing Enterobacterales (CPE), which are resistant to 3^rd^generation cephalosporins and carbapenems, are listed as priority organisms on the WHO Bacterial Priority Pathogens List 2024 [3]. Recently introduced β-lactam/β-lactamase inhibitor combinations, ceftazidime/avibactam or meropenem/vaborbactam, target a range of carbapenemases, but are not active against the metallo-β-lactamase (MBL) family of carbapenemases [4], the most prevalent of which in the UK is *bla*_NDM_ [5].

A recently introduced treatment option for drug resistant Gram negative bacterial infections, including MBLs is cefiderocol, which exploits a ‘Trojan Horse’ mechanism to enter bacterial cells [6]. It mimics bacterial siderophores, molecules that bind extracellular ferric iron that are actively taken up by the bacterial iron acquisition systems, overcoming efflux pumps and porins that reduce periplasmic accumulation of antibiotics [7]. Cefiderocol contains a cephalosporin pharmacophore, which binds to penicillin binding protein 3 in the periplasmic space, inhibiting cell wall synthesis causing cell death [8]. The modified cephalosporin pharmacophore and catechol siderophore chain of cefiderocol reduces hydrolysis, by MBLs making cefiderocol a promising new therapeutic for CPE infection [9].

However, shortly following its introduction, resistance to cefiderocol was reported, and is most frequently encountered in CPE. One retrospective cohort study in England showing that 69.7% of NDM producing Enterobacterales were resistant to cefiderocol based on The European Committee on Antimicrobial Susceptibility Testing (EUCAST) version 14.0 breakpoints, despite no previous exposure to cefiderocol in these patients [10]. Chromosomal mutations associated with cefiderocol resistance are in the iron uptake receptor gene *cirA*, resulting in a non-functional gene, and insertions in the *ftsI* gene encoding penicillin binding protein 3 [11, 12]. However, chromosomal mutations potentially involved in decreased susceptibility to cefiderocol vary by species, with synergistic or antagonistic effects of multiple genomic determinants of cefiderocol resistance not fully understood [11].

Cefiderocol usage in the UK and globally is guided by antibiotic susceptibility testing (AST) in clinical microbiology laboratories, primarily using disk diffusion methodology. EUCAST and the Clinical and Laboratory Standards Institute (CLSI), based in the US, both provide clinical breakpoints to interpret AST results. However, the latest cefiderocol breakpoints in Enterobacterales released by EUCAST and CLSI differ greatly, for both disk diffusion and standard broth microdilution methods [10, 13, 14]. For Enterobacterales isolates that fall in the EUCAST area of technical uncertainty for disc diffusion, other AST methods can be utilised. A gradient cefiderocol strip can be used, but this is validated only for *Pseudomonas* [15], and is not recommended by EUCAST [16]. For other Enterobacterales, broth microdilutions using an iron-depleted cation adjusted Muller Hinton broth (ID-CAMHB) is recommended, although the complexities of this media preparation can cause reproducibility issues [17]. The challenges of conflicting breakpoints together with varying AST methodologies causes variation in susceptibility/resistance classification, raising substantial concerns globally around appropriate cefiderocol usage, antibiotic stewardship and surveillance.

Given that cefiderocol is one of the only treatment options for NDM-producing Enterobacterales, resistance is emerging [18] and there are conflicting AST breakpoints, the aims of our study were two-fold. Firstly, using a clinical isolate collection of NDM producing Enterobacterales from a hospital network in the Northwest of England, we sought to compare AST methods contextualised under both CLSI and EUCAST breakpoints, and secondly, to define the genomic determinants of cefiderocol resistance in these isolates.

## Methods

### Ethics statement

No identifiable data was analysed during this project. As an anonymised use of routinely collected data and following UK Health Research Agency (HRA) guidance the study did not require formal research ethics committee review and was approved by HRA (IRAS ID 345990). Isolates were shared between sites under a Materials Transfer Agreement (ROC10844TE).

## Isolate collection

Isolates were obtained from a previous study in which we characterised an outbreak of NDM producing *Enterobacterales* within a UK hospital network [19]. Fifty-eight NDM producing clinical isolates were initially identified using Xpert Carba-R (Cepheid, US), with the presence of a *bla*_NDM_ gene confirmed by whole genome sequencing (WGS). Clinical isolates were provided by Liverpool Clinical Laboratories on blood agar plates, and a single colony was kept in glycerol stocks at -80°C with storage beads (Micro-Bank, Pro-Lab Diagnostics), to be used within the study. Whole genome sequencing (WGS) data was retrieved from ENA (bioproject PRJEB89345) [19].

## Disk diffusion method for cefiderocol AST

Using the EUCAST disk diffusion method (v.12.0), Millipore NutriSelect® Plus Muller Hinton agar (MHA) (Merek, Germany) was prepared to EUCAST’s guidelines [20] and stored at 4°C before use, and the same batch of agar was used for all tests. Colonies were selected from an overnight LB agar plate and suspended in saline to a 0.5 McFarland standard equivalent density. This suspension was then spread onto the plate in three directions using a swab. A 30µg cefiderocol disk (MAST Group, UK) was placed into the centre of the plate before being incubated at 35°C for 18 hours and measured immediately.

### MIC broth microdilution method for cefiderocol AST of *Escherichia coli* isolates

ID-CAMHB was prepared according to EUCAST and CLSI guidance, always using the same batch of autoclaved Millipore NutriSelectCAMHB (Merck, Germany) [21, 22]. Cefiderocol (Shionogi, Japan) was dissolved in DMSO to a final concentration of 2mg/mL with single use aliquots stored at -20°C.

*E. coli* inoculum was prepared by passaging a single colony from an overnight MHA plate in 10 mL ID-CAMHB for 8-18 hours, 200rpm at 37°C. This was diluted to an OD600nm of 0.001 (corresponding to 10^5^ cfu/mL) in ID-CAMHB. To prepare MIC plates, cefiderocol was serially diluted across a nine-fold dilution range in 96-well round bottom microtiter plates (0.5mg/L to 128mg/L) in ID-CAMHB in triplicate wells per drug concentration. Equal volume of 50 µL of *E. coli* suspension was added to wells containing serial diluted cefiderocol, including drug free wells as positive growth controls, giving a test range of 0.25mg/L to 64mg/L. Inoculum free ID-CAMHB were included as media contamination controls. *E. coli* ATCC 25922 was included alongside all *E. coli* isolates as a quality control reference as recommended by EUCAST and CLSI [22, 23]. Microtiter plates were incubated statically at 35°C for 16-20 hours and read immediately using guidance documents from EUCAST and CLSI, where at least two wells in agreement were taken as the MIC [21, 22]. MICs were determined a minimum of three times independently.

### MIC gradient strip method for cefiderocol AST

Following the protocol for MIC gradient strip testing as provided by the manufacturer, colonies from an overnight LB agar plate were suspended in saline to a density equal to a 0.5 McFarland standard [24]. This suspension was then swabbed onto an 25mL MHA pre-poured media plate (E&O Labs, United Kingdom) in three directions. An MIC gradient strip (Liofilchem, Italy) was placed into the centre of the plate before being incubated at 35°C for 18 hours and read immediately.

### Analysis and comparison of disc diffusion, MIC gradient strip and microbroth dilution AST

Both EUCAST (v.15.0, [25]) and CLSI (2023 breakpoints, [26]) breakpoints were used to determine susceptibility to cefiderocol for disk diffusion and broth microdilution. Both sets of breakpoints were experimentally applied to the gradient strip data in order to make a comparison between the gradient strip and other AST methods. MIC_50_ and MIC_90_ were calculated from broth microdilution using the AMR package for R [27]. When comparing different AST methods, category agreement is defined as an isolate generating the same category (susceptible, intermediate, resistant), while essential agreement is defined as an isolate producing an MIC that is ± one doubling dilution concentration. When evaluating category agreements, a minor error is when an isolate was susceptible versus intermediate or intermediate versus resistant between different methods. A major error occurs where resistance was identified by an AST method (disc diffusion, MIC gradient strip), when the reference standard AST method (microbroth dilution) identified susceptibility. Finally, a very major error occurs when an AST method (disc diffusion, MIC gradient strip) identified an isolate as susceptible, but microbroth dilution identifies resistance [28]. For disc diffusion, isolates assigned the ATU category were excluded from category agreement calculations as these could not be assigned a category of susceptible, intermediate or resistant [29].

When determining category agreement for sensitive, intermediate or resistant *E. coli* isolates for disc diffusion and MIC gradient test strip, the broth microdilution was used as the reference standard. When calculating category agreement between disk diffusion and MIC gradient test strip methods on the full isolate set, the disk diffusion method was used as the reference methodology to measure disagreements. The gradient MIC test strip is only validated for use in *Pseudomonas* and so was not used as a reference method in this study.

### Identification of β-lactamase genes and genetic mutations implicated in cefiderocol resistance

A literature search was conducted using Pubmed to identify previously reported cefiderocol resistance genomic determinants to produce a bespoke ARIBA (v.2.14.6) [30] database per genus for *Citrobacter, Escherichia, Enterobacter*, and *Klebsiella*, reflecting the isolates in our study. This search provided no results for chromosomal genes linked to cefiderocol resistance for the *Citrobacter* spp. (last day of retrieval, 23^rd^ July 2024) so relevant chromosomal genes for other species were included in the *Citrobacter* spp. ARIBA database. β-lactamase genes included from the literature as being linked to cefiderocol resistance were *bla*_SHV_; *bla*_CTX-M_; *bla*_CMY_; carbapenemase genes.

Twelve reference genomes were selected for each genus, including at least one reference genome from an isolate previously described as susceptible to cefiderocol (Table S2-Table S5). The sequences of the chromosomal genes of interest were obtained through the NCBI Prokaryotic Genome Annotation Pipeline [31]. Clustal omega (v.1.2.4) [32] was used to align the protein sequences of chromosomal genes of interest from these reference genomes and identify the wild-type sequence for each chromosomal gene [32]. Based on this, a singular reference genome was then selected, and the relevant gene sequences extracted, to build a bespoke ARIBA database for each genus.

ARIBA searches were conducted for each genus across three databases; custom, SRST2 and CARD. Selected β-lactamase genes were identified using the SRST2-ARGANNOT [33] database as implemented in ARIBA (downloaded 2nd May 2024) [33]. Mutations within chromosomal genes of interest were identified using the genus specific bespoke ARIBA database. For *K. pneumoniae* isolates, the *ompK35, ompK36*, and *ompK37* genes were added to the search using the CARD [34] database as implemented in ARIBA (downloaded 14^th^ February 2024) [34].

## Results

The collection of isolates within this study included *Enterobacter hormachei* (n=20), *Klebsiella pneumoniae* (n=20), *Escherichia coli* (n=14), and *Citrobacter freundii* (n=3). One isolate was identified to contain additional carbapenemase genes using Gene Xpert Carba-R at the point of isolation, with *bla*_OXA-48 like_ identified (Table 1).

**Table 1.** Characterisation of the isolates in this study and their carbapenemase genes identified using WGS.

| Species | Carbapenemase genes | Isolates (n) |
| --- | --- | --- |
| *Citrobacter freundii* | NDM | 3 |
| *Enterobacter hormaechei* | NDM | 20 |
| *Escherichia coli* | NDM | 13 |
| *Escherichia coli* | NDM + OXA-48 | 1 |
| *Klebsiella pneumoniae* | NDM | 20 |

### Discrepancy of disk diffusion breakpoints has significant impact on resistance assessment

The disk diffusion method, based on EUCAST interpretation, classified 94.7% (n=54/57) of the isolates as cefiderocol resistant and 5.3% as susceptible (n=3/57). However, 22.8% (n=13/57) were within the Area of Technical Uncertainty (ATU) (Figure 1).

**Figure 1:**
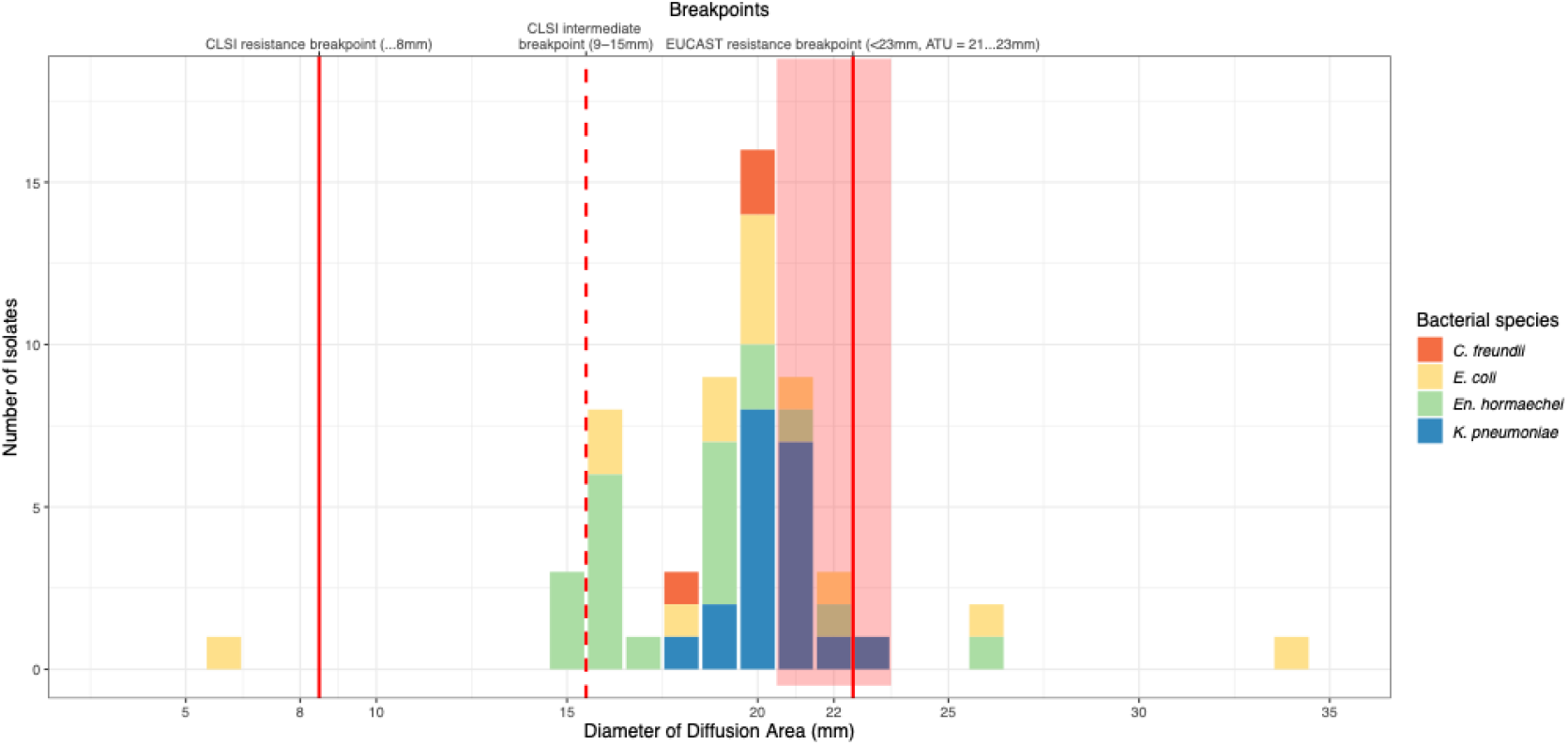
Cefiderocol disk diffusion diameters of collection isolates considering CLSI and EUCAST breakpoints. Counts of isolates per disk diffusion area of all isolates studied, colour indicates species as indicated in the legend. Red lines indicate CLSI and EUCAST resistance breakpoints, dashed line indicates CLSI intermediate breakpoint, shaded red area indicates EUCAST area of technical uncertainty (ATU).

Approximately two thirds of *E. coli* (71.4%, n=10/14) and over three quarters of *En. hormaechei* (85%, n=17/20) isolates were classified resistant to cefiderocol, and outside of the ATU. Of these isolates only one *E. coli* was also classified as resistant using the CLSI breakpoint (1/57, 1.8%), and additionaly three *En. hormaechei* were classified as intermediate (3/57, 5.26%).

### Disc diffusion overestimates cefiderocol resistance compared to broth microdilution using EUCAST breakpoint

To assess resistance classification agreement between different methods, we tested resistance classification using broth microdilution, the reference standard in AST for Enterobacteriaceae [35], with all *E. coli* isolates. Three biological repeats were performed for all isolates, however given reproducibility issues encountered when combining results from different batches of iron (Fe)-depleted media preparations, four isolates required a total of six biological repeats to achieve three results with essential agreement. The results of the ATCC 25922 quality control strain were in essential agreement in all microbroth dilution experiments. Using the modal average of all MIC results derived from our isolates, the MIC_50_ was 1mg/L and the MIC_90_ was 6.8mg/L; the range of MIC values obtained were 0.5mg/L to >64mg/L (Figure 2), and 21.4% were classified resistant to cefiderocol using the EUCAST breakpoint (n=3/14). There was a 70.3% decrease in the number of isolates initially identified as resistant by disc diffusion (n=10/14). None of the isolates within the ATU for disk diffusion were classed as resistant according to broth microdilution, suggesting disc diffusion over-estimates cefiderocol resistance. Applying CLSI breakpoints for broth microdilution results, one isolate (NDM138) classified intermediate (7.1%, n=1/14) and one isolate (NDM150) resistant (7.1%, n=1/14) (Figure 2).

**Figure 2.**
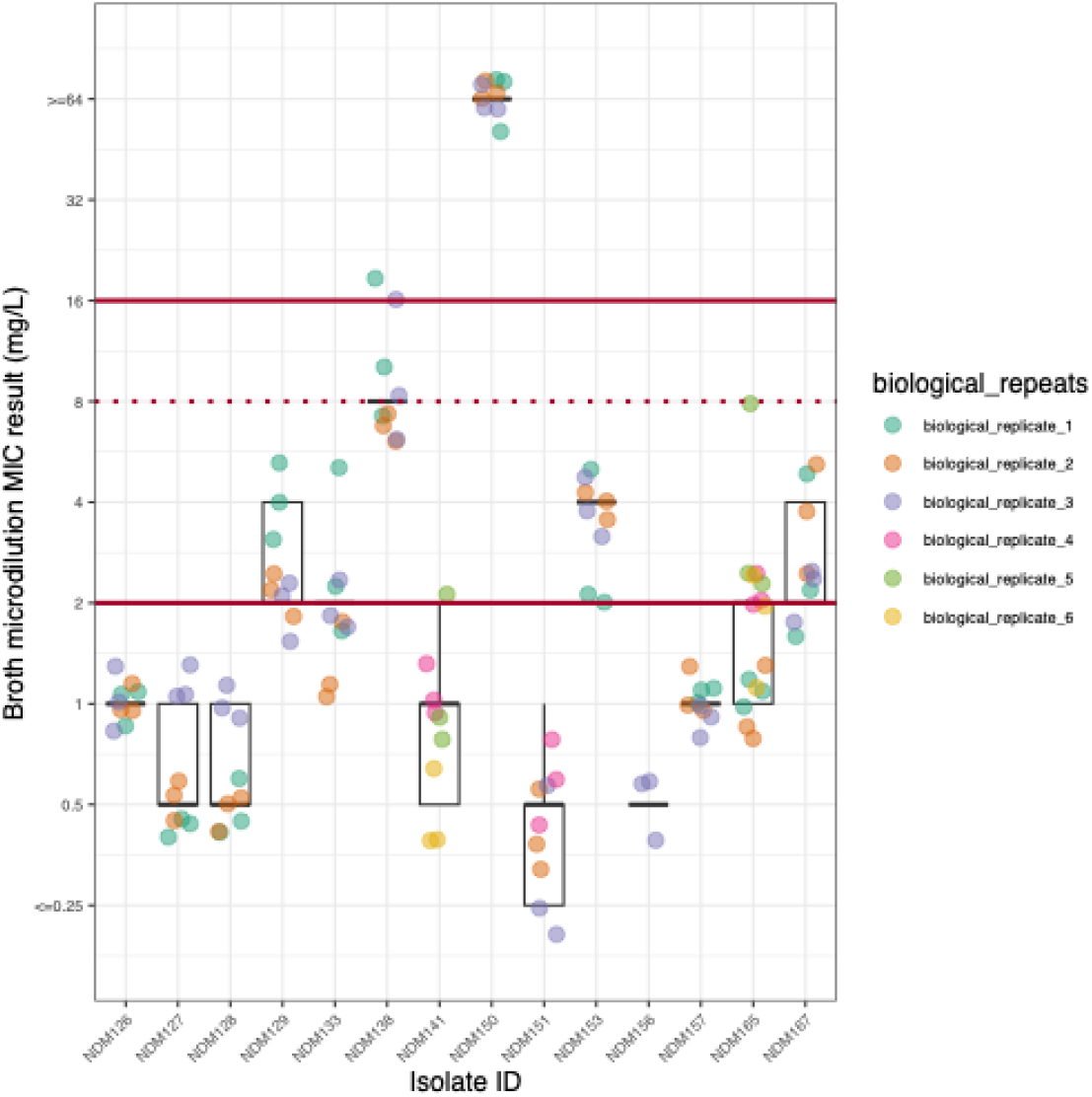
Variation of cefiderocol MIC values by broth microdilution assessing biological and technical replicates. Boxplots showing different broth microdilution MIC results per isolates of six biological replicates as indicated by colours, and three technical replicates per biological replicate.

The category agreement for *E. coli* isolates between broth microdilution and disk diffusion was 38.5% applying EUCAST breakpoints (excluding ATU isolates, n=5/13) with major errors for eight isolates when using disc diffusion (61.5%, n=8/13). Using CLSI breakpoints, the category agreement was 92.9% (n=13/14); with one instance of a minor error (Figure 3A; Figure 3B).

**Figure 3.**
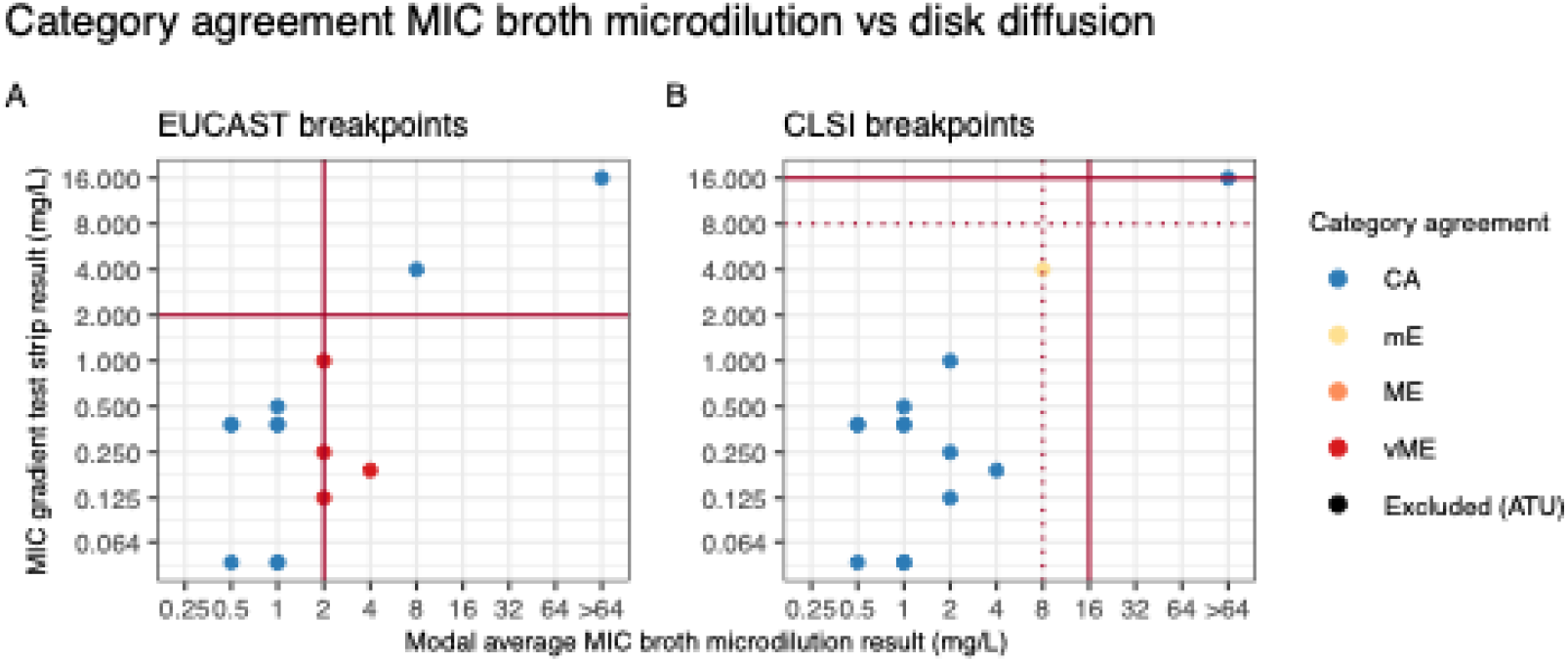
Category agreement of MIC broth microdilution vs disk diffusion. We show modal average of our MIC broth microdilution experiments against MIC gradient test strip result under EUCAST (A) or CLSI (B) breakpoints; indicating category agreement in the colour scale with Categorical Agreement (CA) in blue, minor Error (mE) in yellow, Major Error (ME) in orange, very Major Error (vME) in red, and excluded values (ATU) in black.

### Higher category agreement of AST methods using CLSI vs EUCAST breakpoints potentially driven by categorical underestimation of resistance with CLSI

In addition to disc diffusion category agreement with broth microdilution, a MIC gradient test strip was also tested and compared to disc diffusion (n=57) and broth microdilution (n=14). Using MIC gradient test strip data, the MIC_50_ for all isolates was 0.25mg/L and the MIC_90_ was 3mg/L; the range of MIC values obtained were 0.047mg/L to 16mg/L.

With the MIC gradient test strip, the EUCAST breakpoint classified 12.3% (n=7/57) as resistant, whereas the CLSI breakpoint classified one isolate (1.75%) as resistant and 5.26% isolates (n=3/57) as intermediate. The category agreement between the MIC gradient test strip and disk diffusion method was 22.7% when using the EUCAST breakpoint (excluding 13 ATU isolates, n=10/44), and 96.5% when using the CLSI breakpoint (n=55/57) (Figure 4).

**Figure 4.**
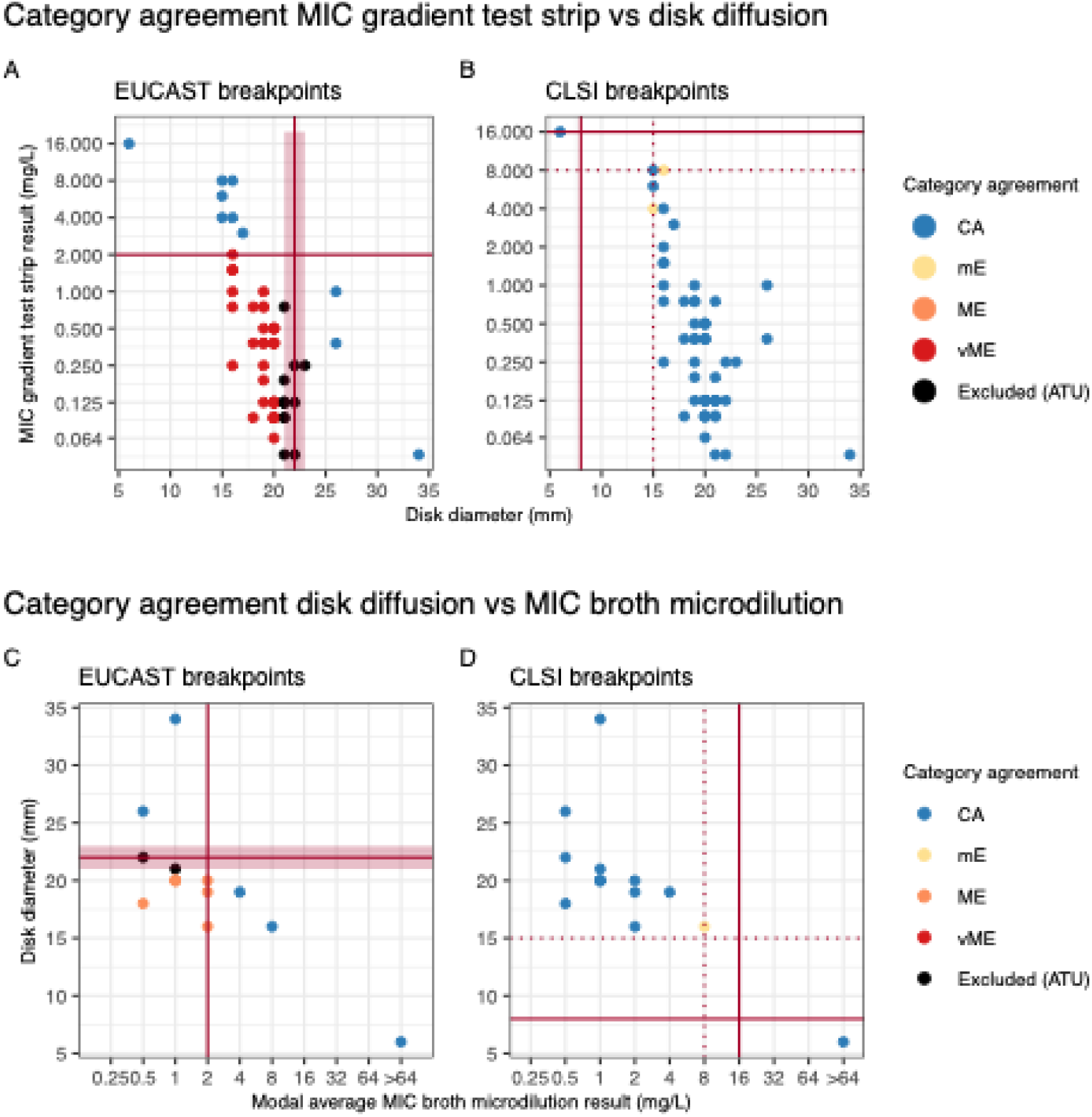
Comparing categorical agreement of different MIC measurements. We show category agreement of disk diffusion vs MIC gradient test strip for EUCAST (A) and CLSI (B) breakpoints; and modal average of the MIC broth microdilution results against disk diffusion under EUCAST (C) and CLSI (D) breakpoints. Colours indicate category agreements as in Figure 5.

The category agreement between MIC gradient test strip and broth dilution was 71.4% applying EUCAST breakpoints (n=10/14), with four cases of very major errors (28.6%, n=4/14). There was 92.9% category agreement using CLSI breakpoints (n=13/14), with one minor error recorded (Figure 4C; Figure 4D). The essential agreement between broth microdilution and the MIC gradient test strip in these *E. coli* isolates was 35.7% (n=5/14).

### Identification of beta-lactamase genes and putative determinants of cefiderocol resistance

WGS identified *bla*_NDM-1_ (74.1%, n=43/57), *bla*_NDM-5_ (20.7%, n=12/57), *bla*_NDM-6_ (n=1), and *bla*_NDM-8_ (n=1) in our isolates, and *bla*_OXA-48-like_ in a single isolate (Table 1). Mutations within chromosomal genes were identified in all species (FigS2-S4).

Of the *E. coli* with *bla*_NDM-5_ (10/14), 9/10 of these isolates were categorised as resistant using EUCAST interpretation of the disk diffusion results. Of the isolates with *bla*_NDM-1_ (4/14), 1/4 was susceptible, 2/4 were ATU and 1/4 resistant. Several isolates co-carried another beta-lactamase gene including *bla*_CTX-M-15_ (6/6 resistant), *bla*_CMY-42_ (2/2 resistant), *bla*_SHV-12_ (2/3 ATU, 1/3 susceptible), and *bla*_OXA-48-like_ (1/1 resistant) (Figure 5A,B, S2). All *E. coli* isolates had mutations identified in at least two of the chromosomal genes analysed (*cirA, ftsI, fiu, tonB)*. The two isolates with the highest MIC had the combination of *bla*_NDM-5_, *bla*_CMY42_, and were the only two *E. coli* isolates with the same frameshift mutation in *cirA* (Ser-90) (Figure 5A,B, S2).

**Figure 5.**
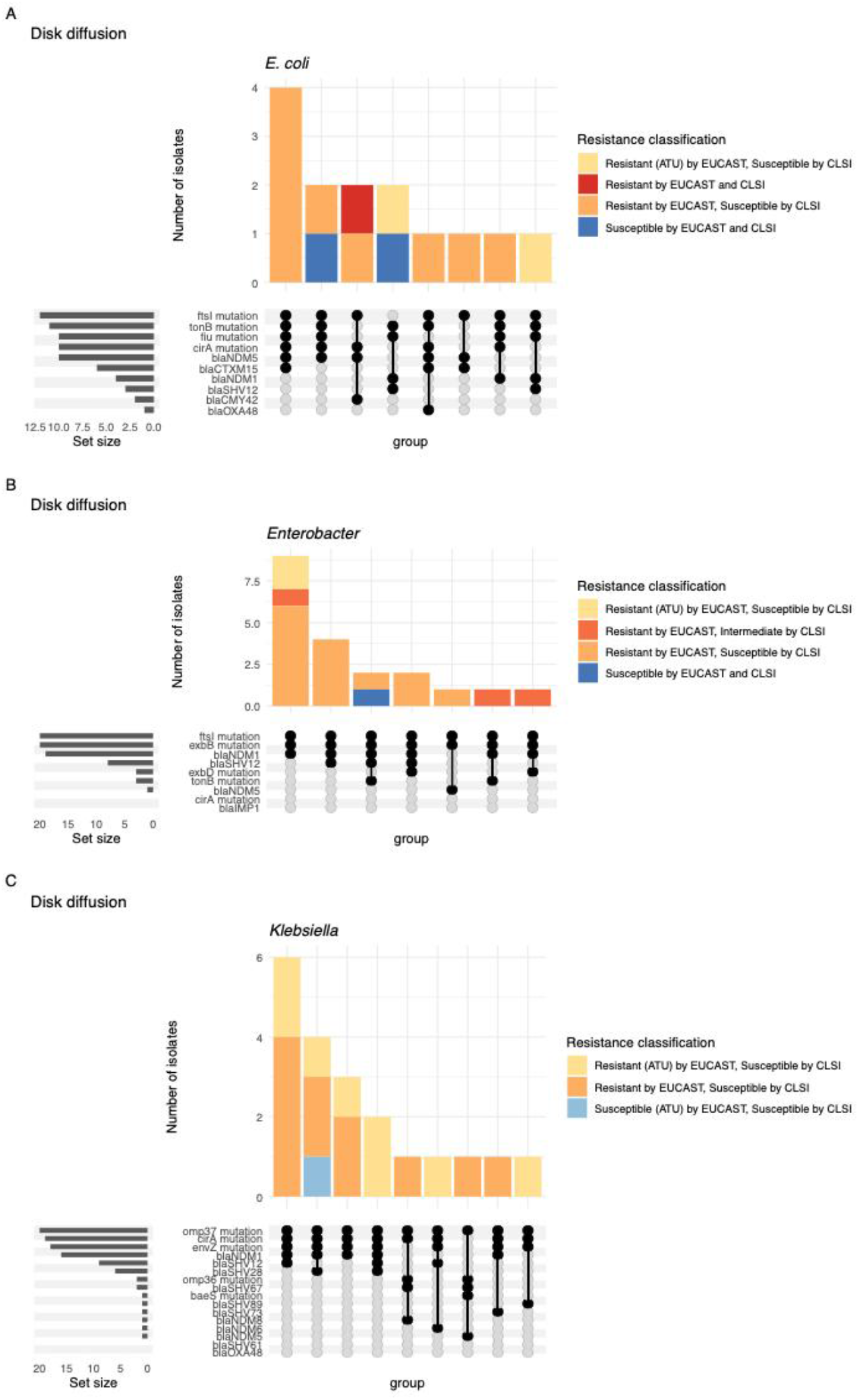
Presence of mutations previously described of relevance for cefiderocol resistance in the genomic data of our study isolates. Upset plots showing the number of isolates for each combination of relevant mutations for *E. coli* (A), *Enterobacter* (B) and *Klebsiella* (C). Resistance classification according to disk diffusion for EUCAST and CLSI are as indicated in the colour legends.

Nineteen of the *Enterobacter* isolates carried the *bla*_NDM-1_ gene (1/19 susceptible, 2/19 ATU, 16/19 resistant using the EUCAST breakpoint). The gene *bla*_NDM-5_ occurred in a single isolate, which was resistant. *bla*_SHV-12_ was found in eight isolates (1/8 susceptible, 7/8 resistant) (Figure 5C, S3). No *cirA* gene was identified in *En. hormachei* and the *tonB* and *wxbD* genes were absent in most isolates (n=18/22 and n=16/22, respectively). All *Enterobacter* isolates harboured at least one mutation in the *ftsI* gene, and the previously unreported R587V mutation was identified in the three isolates with the smallest disk diffusion diameter of the group (Figure 5C, S3).

Sixteen of the *Klebsiella* spp. encoded *bla*_NDM-1_ (1/16 susceptible, 6/16 ATU, 9 resistant), with *bla*_NDM-5_ (1/1 resistant), *bla*_NDM-6_ (1/1 ATU) and *bla*_NDM-8_ (1/1 resistant) each detected in a single isolate (Figure 5D, S4). All co-carried *bla*_SHV_ genes, including *bla*_SHV-12_ (9/16), *bla*_SHV-28_ (4/16), *bla*_SHV-67_ (1/16), *bla*_SHV-73_ (1/16) and *bla*_SHV-89_ (1/16) and had *ompk37* mutations, and 17/19 had mutations in *cirA* and *envZ*. The two isolates without mutation in *cirA* and *envZ* had mutations in *ompk36*, and both were resistant according to EUCAST disk diffusion breakpoints. Across all species the mutations in the genes associated with cefiderocol resistance were diverse (Figure 5D, S4); for example in the *E. coli* strains alone fourteen different amino acid changes in *cirA* were recorded.

## Discussion

Appropriate usage of cefiderocol to treat CPE infection relies on the accuracy of AST, to identify patients with susceptible infections. We found that the discrepancy between EUCAST or CSLI breakpoints had a major impact on the proportion of isolates deemed susceptible to the antibiotic, with the majority of isolates being classified as resistant by EUCAST but susceptible by CSLI. The EUCAST breakpoint was developed using a collection of Enterobacterales including a higher proportion of resistant isolates than in the studies that informed the CSLI breakpoint, resulting in the more conservative breakpoint values for BMD and DD.

Whilst EUCAST breakpoints are widely used in Europe, and CSLI in the United States, the selection of either breakpoint varies globally, and both are endorsed for use by the World Health Organization’s Global Antimicrobial Resistance Surveillance System (GLASS) [36]. This major discrepancy in the cefiderocol breakpoint between the two bodies will lead to major impacts on resistance surveillance and epidemiology globally, and differences in how cefiderocol is used clinically. This has previously been highlighted for other pathogen-drug combinations which have different CLSI/EUCAST breakpoints, eg ciprofloxacin-Enterobacterales [37].

The majority of AST performed clinically for any antibiotic is the DD method. Technical complexities and time cost of broth microdilution at the throughput necessary for hospital microbiology laboratories make it challenging to implement routinely, and the need for iron depleted media makes this process even more onerous for cefiderocol. For the *E. coli* isolates tested with both methods we found poor agreement between DD and broth microdilution with EUCAST breakpoints, with the commonly used DD method overestimating the number of isolates resistant, considering the broth microdilution as the reference standard. If representative of cefiderocol AST more broadly, this has the potential to overestimate resistance, meaning cefiderocol therapy is discounted in cases where it would have been an effective antibiotic. Greater category agreement was found when applying the CSLI method, driven by the characterisation of only a single isolate as resistant using DD or broth microdilution using this less conservative breakpoint.

Implementation of broth microdilution in a clinical setting remains challenging, and in the case of cefiderocol reproducibility issues originating from the brand of media utilised need to be accounted for [38, 39]. However, commercial *in vitro* AST test kits are available, such as comASP cefiderocol (liofilchem, Italy) and the Sensititre™ Gram Negative RUO Susceptibility Testing Plate (Thermo Fischer, US), which may provide a means for performing reference standard AST in a manner implementable by clinical microbiology services [40], however a EUCAST warning against such commercially available tests remains in place [16].

In addition to technical challenges, current use is mainly in patients with complicated clinical history, making it challenging to define the effect of cefiderocol itself in the context of complex treatment history. There is thus limited insight on treatment success or failures based on *in vitro* resistance profile, as cefiderocol use often follows or is in parallel with several other antimicrobials and in patients with severe co-morbidities.

The molecular mechanisms of cefiderocol resistance in Enterobacterales is complex, with reduced susceptibility typically a combination of carbapenemase production, multiple beta-lactamases, and chromosomal mutations. Mutations in the catecholate receptor *cirA*, causing a deficiency in the siderophore transport system and therefore reducing cefiderocol uptake, have a major impact on cefiderocol susceptibility, and were identified in the isolates with the smallest DD zone diameters in this study. Frameshift mutations, causing a non-functional protein were found in the isolates with the most reduced susceptibility. However not all mutations in *cirA* are predictive of cefiderocol resistance, as amino acid changes were identified in several susceptible isolates. Mutations in other siderophore receptor genes, including *exbD, fiu* and *tonB* were also identified.

We identified several resistant isolates with mutations in the *ftsl* gene encoding penicillin binding protein 3, including those (A413V, Q227H, E349K and I532L) linked to decreased susceptibility to aztreonam-avibactam [41]. We also detected the plasmid-mediated AmpC gene *bla*CMY-42, also linked to resistance to aztreonam-avibactam. This raises the possibility of cross resistance to both therapeutics emerging in NDM producing isolates, which is concerning given these are both part of the limited available treatments for these organisms.

Predicting resistance and assessing resistance profiles is further complicated by the genomic plasticity of Enterobacteriaceae. The *cirA* gene is absent in *Enterobacte*r isolates as reported previously [42], and *ftsl* mutations are widespread in *E. coli* and *Enterobacter* but not in *Klebsiella*. Phenotypic differences were also noted, with smaller average DD zones found in the *Enterobacter hormaechei* isolates. This highlights the difficulties inferring cefiderocol resistance from sequencing data and interpretation of AST with possible differences in phenotypic readout. Genomic plasticity and differences in resistance determinants further complicates the design of a molecular diagnostic assays to identify potentially resistant isolates to guide treatment, again underscoring the importance of accurate phenotypic assays and suitable breakpoints.

Our study has several limitations. This study occurred in a hospital network in a single city in the UK, and the isolates and MIC ranges determined here may not be representative of other settings. Broth microdilution was solely tested on the *E. coli* isolates, due to time and resource constraints, only allowing the assessment of the agreement between the BMD and DD methodologies with this species. The MIC test strip is approved for *Pseudomonas* and not for Enterobacterales, and is not recommended for clinical AST, however we included it within the study as a commercially available potential methodology, in order to assess its agreement to the clinically used methods. Whilst we used whole genome sequencing to characterise the various mutations and genes present in the isolates with known links to cefiderocol resistance, the study was not powered to assess their relative contributions to cefiderocol MIC, or identify resistance determinants that were not previously reported. The relationship between cefiderocol MIC and the likelihood of clinical failure is currently unknown, and further *in vitro* and *in vivo* pre-clinical studies and clinical trials are needed to robustly address this research gap.

This study highlights the impact of the selection of CLSI or EUCAST breakpoints on cefiderocol AST interpretation, and on the category agreement of various AST methods. The mechanism of cefiderocol resistance in Enterobacterales is complex and differs between species, and it is unclear whether this leads to different phenotypic baseline readout that might necessitate different breakpoints between different species within the Enterobacterales to ensure clinically appropriate use.

## Funding information

EH acknowledges funding from Wellcome (217303/Z/19/Z) and the BBSRC (BB/V011278/2). TE acknowledges funding from an Academy of Medical Sciences Springboard award (REF: SBF009\1181). DC acknowledges funding from Innovate UK Engineering Biology award (10073149).

## Author contributions

Conceptualization: DL, TN, JC, JML, AJF, CB, EH, TE. Data curation: DL, TN, JC, VO, JML, CB. Formal analysis: CD, DC, AJF. Funding acquisition: DC, EH, TE. Investigation: CD, DC, DL, TN, JC, VO. Methodology: DC, FEG, EH, TE. Project administration: JML, CB, EH, TE. Software: CD, AJF, EH. Resources: DL, TN, JC, VO, FEG, JML, CB. Supervision: JML, CB, EH, TE. Validation: CD, DC, TN, JC, FEG, JML, CB. Visualization: CD, EH. Writing – original draft: CD, DC, EH, TE. Writing – review & editing: all authors

## Conflict of interest

None to declare.

## Supplementary figures

**Figure S1.**
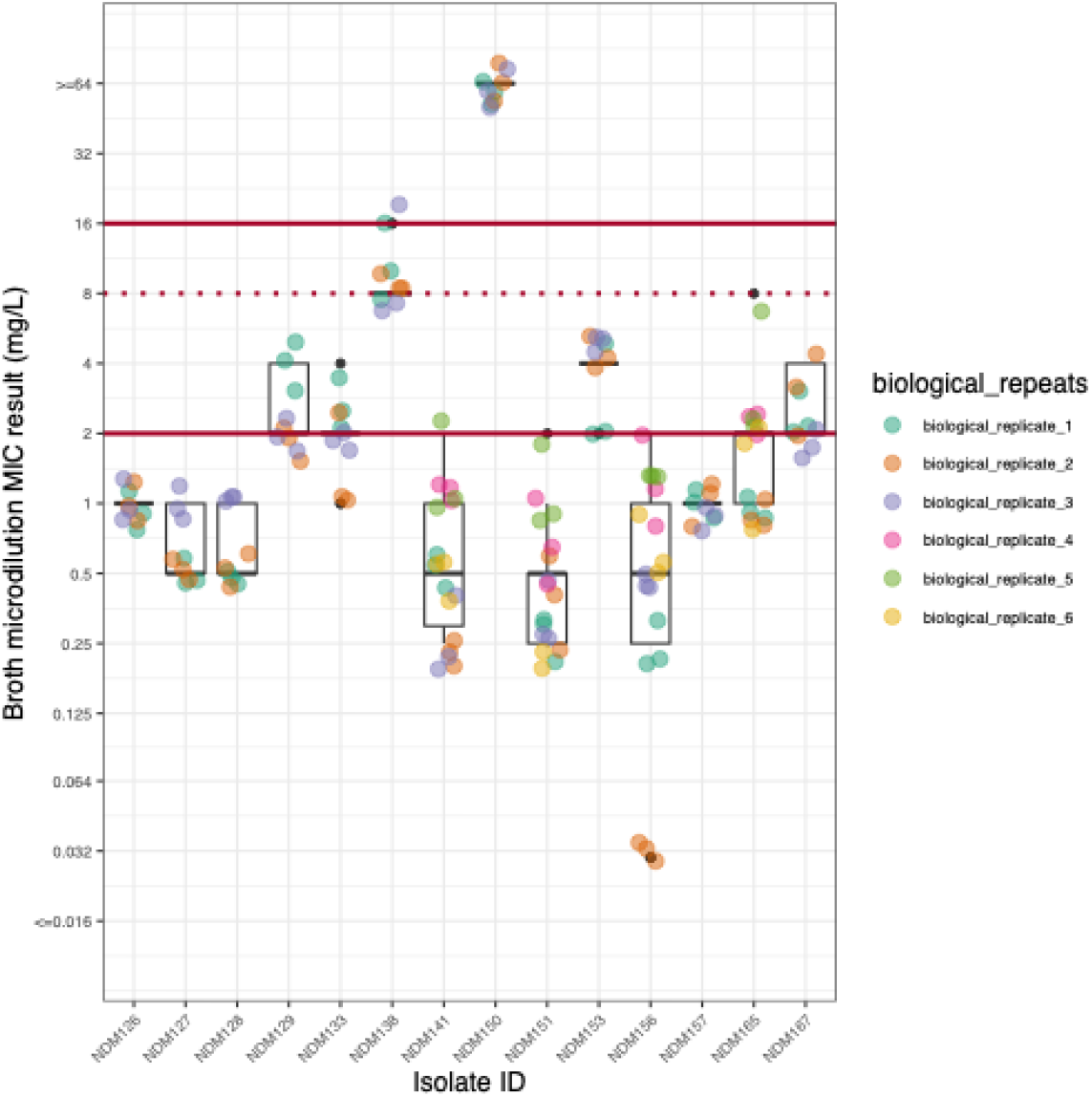
Broth microdilution results for all replicates. Broth microdilution results for all replicates of the *E. coli* isolates in our study.

**Figure S2.**
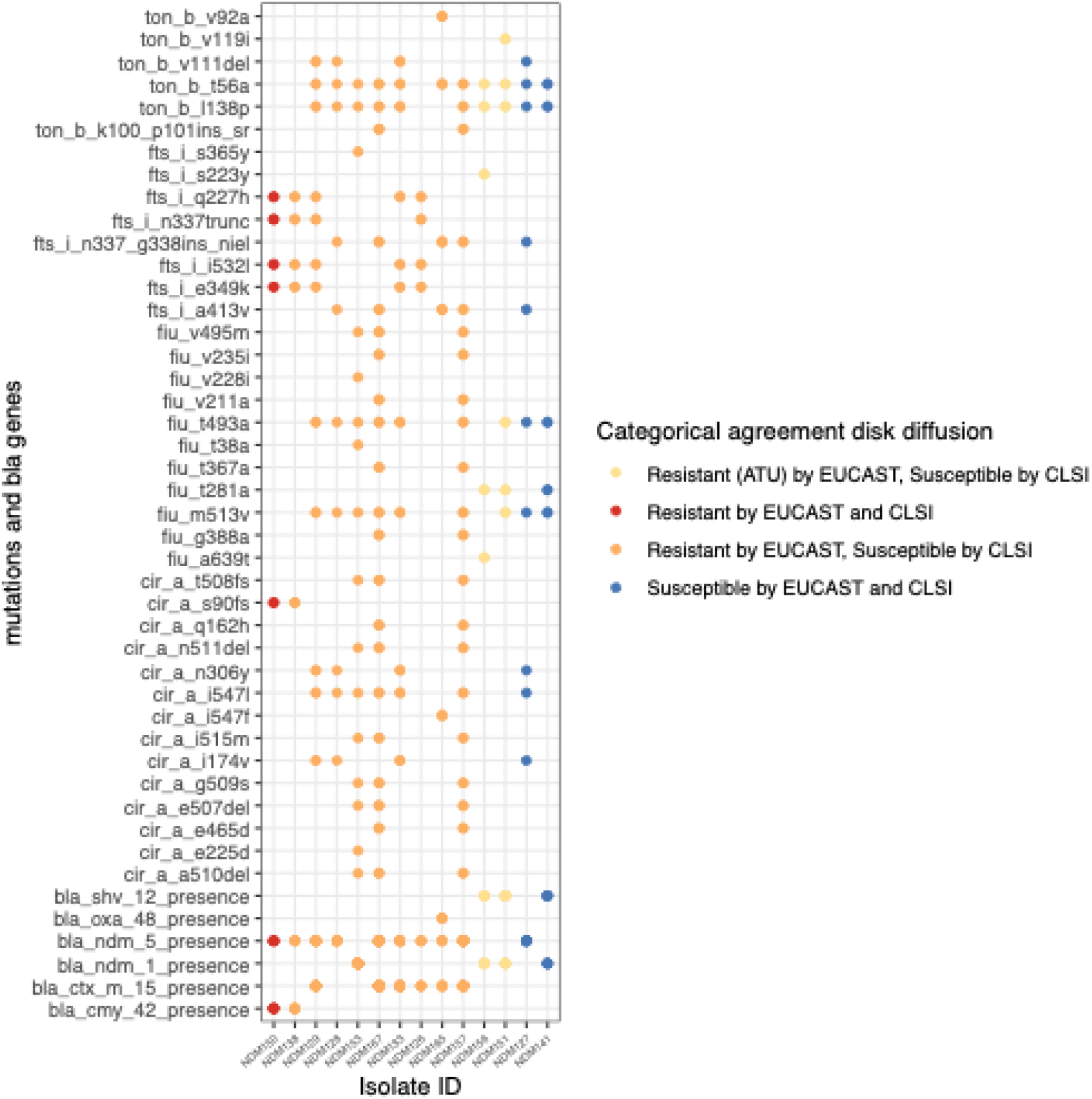
Bubble plot diagram indicating presence/absence of mutations and carbapenemase genes (rows) for the *E. coli* isolates in our collection (columns). Categorical agreement of disk diffusion values as indicated in the colour legend.

**Figure S3.**
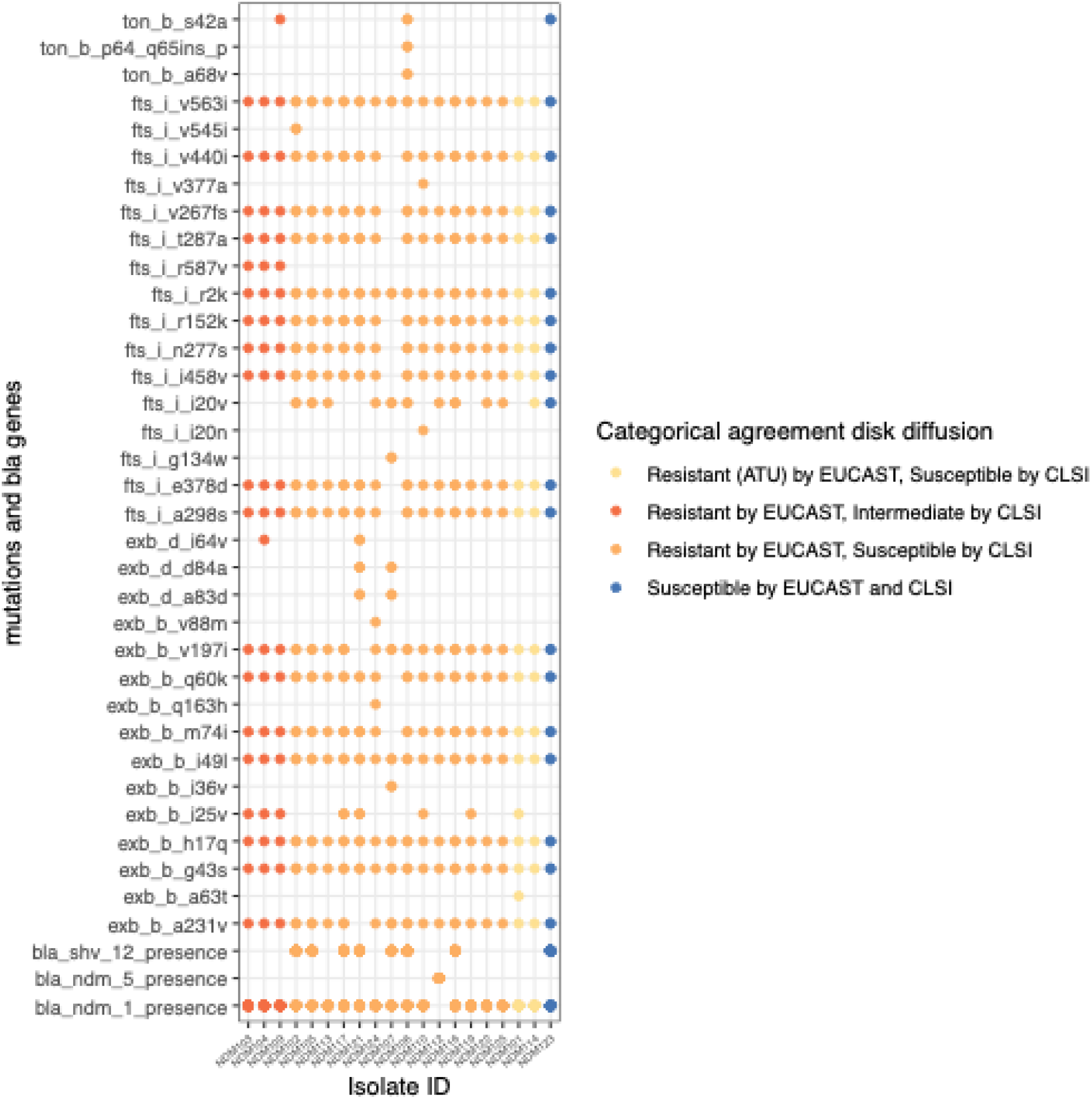
Bubble plot diagram indicating presence/absence of mutations and carbapenemase genes (rows) for the *Enterobacter* isolates in our collection (columns). Categorical agreement of disk diffusion values as indicated in the colour legend.

**Figure S4.**
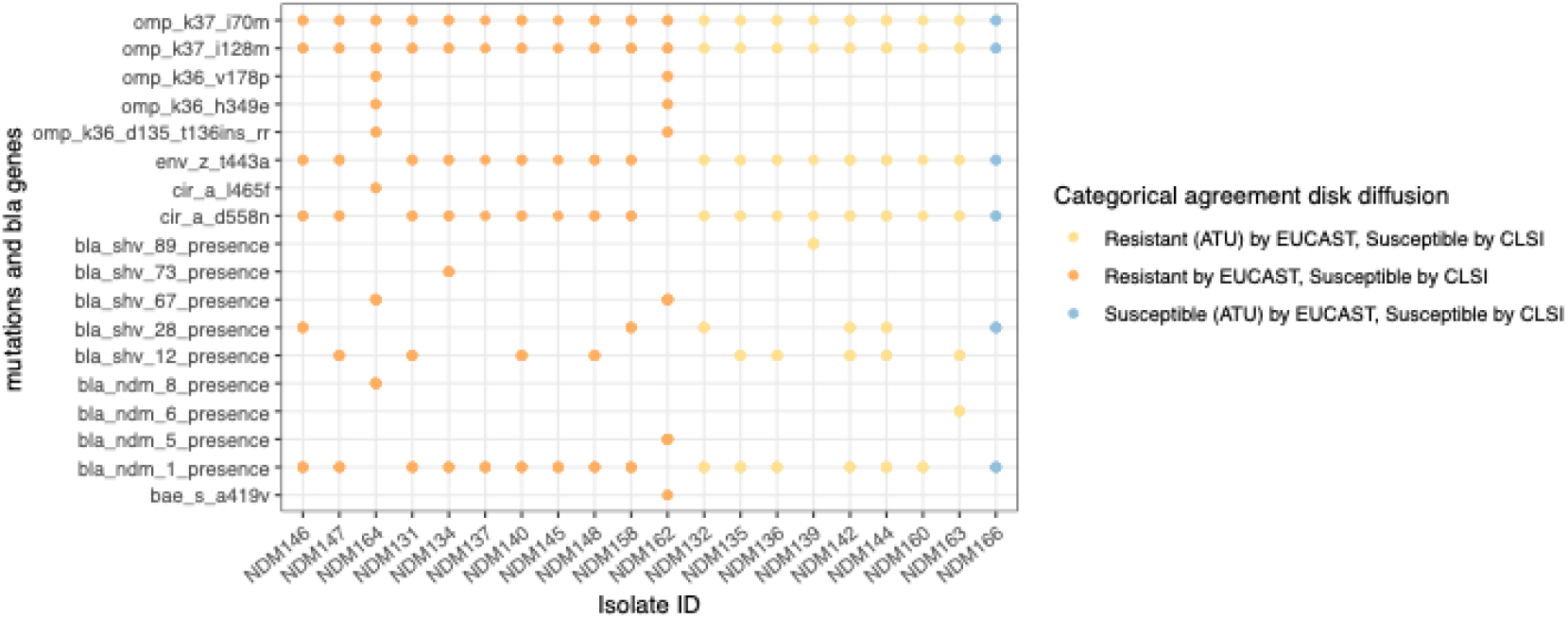
Bubble plot diagram indicating presence/absence of mutations and carbapenemase genes (rows) for the *Klebsiella* isolates in our collection (columns). Categorical agreement of disk diffusion values as indicated in the colour legend.

## Notes

### Competing Interest Statement

The authors have declared no competing interest.

